# Patient iPSC models reveal glia-intrinsic phenotypes in multiple sclerosis

**DOI:** 10.1101/2023.08.01.551553

**Authors:** Benjamin L.L. Clayton, Lilianne Barbar, Maria Sapar, Tomasz Rusielewicz, Kriti Kalpana, Bianca Migliori, The NYSCF Global Stem Cell Array® Team, Daniel Paull, Katie Brenner, Dorota Moroziewicz, Ilana Katz Sand, Patrizia Casaccia, Paul J. Tesar, Valentina Fossati

**Affiliations:** Department of Genetics and Genome Sciences, Case Western Reserve University School of Medicine, Cleveland, OH, 44106, USA; The New York Stem Cell Foundation Research Institute, New York, NY 10019, USA; Current affiliation: Department of Developmental Biology, Washington University School of Medicine, St. Louis, MO, 63105, USA; Corinne Goldsmith Dickinson Center for Multiple Sclerosis, Department of Neurology, Icahn School of Medicine at Mount Sinai, New York, NY 10129, USA; Advanced Science Research Center at CUNY, New York, NY 10031, USA

## Abstract

Multiple sclerosis (MS) is considered an inflammatory and neurodegenerative disease of the central nervous system, typically resulting in significant neurological disability that worsens over time. While considerable progress has been made in defining the immune system’s role in MS pathophysiology, the contribution of intrinsic CNS-cell dysfunction remains unclear. Here, we generated the largest reported collection of iPSC lines from people with MS spanning diverse clinical subtypes and differentiated them into glia-enriched cultures. Using single-cell transcriptomic profiling, we observed several distinguishing characteristics of MS cultures pointing to glia-intrinsic disease mechanisms. We found that iPSC-derived cultures from people with primary progressive MS contained fewer oligodendrocytes. Moreover, iPSC-oligodendrocyte lineage cells and astrocytes from people with MS showed increased expression of immune and inflammatory genes that match those of glial cells from MS postmortem brains. Thus, iPSC-derived MS models provide a unique platform for dissecting glial contributions to disease phenotypes independent of the peripheral immune system and identify potential glia-specific targets for therapeutic intervention.

## Introduction

Multiple sclerosis (MS) is a chronic inflammatory and neurodegenerative disease of the central nervous system (CNS), and the leading cause of non-traumatic neurological disability in young adults^1^. MS is a multifactorial disease resulting from a complex interplay between environmental risk factors (such as EBV infection^2^) and genetic predisposition^3,4^. During the early phase of the disease, MS most commonly manifests with focal neurological symptoms caused by acute demyelinating lesions, with some degree of endogenous repair (relapsing remitting MS, RRMS^5^). Some individuals with MS (less than 15%) experience a progressive course from disease onset that is associated with a worse prognosis (primary progressive MS, PPMS^6^). Over time, most RRMS individuals will exhibit a secondary progressive phenotype, characterized by steady accumulation of neurological disability related to failure of repair mechanisms and consequent neurodegeneration (secondary progressive MS, SPMS)^7^. In RRMS, infiltration of peripheral immune cells and inflammation predominate and disease-modifying therapies targeting B or T cells dramatically reduce the development of new lesions and relapses^8,9^. Unfortunately, these therapies are at best modestly effective at preventing the neurodegeneration that drives disease progression^10^ suggesting disease mechanisms that are independent of peripheral immunity. The pathogenic mechanisms that drive chronic progression in MS are only partially understood, highlighting an urgent unmet need.

Accumulating single-cell transcriptome profiles of postmortem brains have increased our understanding of glial-specific changes in MS^11–13^, involving astrocytes and oligodendrocytes. In addition, GWAS and epigenomic studies have highlighted glial aberrations in people with MS that are independent of the peripheral immune system^14,15^. Studies of postmortem brains, however, cannot discern the intrinsic phenotypes of glial cells in MS from the effects of inflammatory stimuli and peripheral immune cells. Human models based on induced pluripotent stem cell (iPSC) technology serve as important systems to investigate complex, multifactorial diseases. In particular, patient iPSC-based studies are used extensively to model CNS disorders that cannot be fully recapitulated by animal models^16,17^. iPSC modeling therefore provides the opportunity to investigate glial cell dysfunction in MS.

Here, we combined the power of iPSC disease modeling and single-cell transcriptomics to identify glial cell-intrinsic MS phenotypes that occur in the absence of inflammatory stimuli or interactions with peripheral immune cells. We found that iPSC-derived cultures from people with primary progressive MS generated fewer oligodendrocytes. Moreover, iPSC-oligodendrocyte lineage cells and iPSC-astrocytes from people with MS showed increased expression of immune and inflammatory genes that match those of cells from MS post-mortem brains. This study highlights the value of iPSC modeling for generating disease-relevant cell types and capturing glial cell-intrinsic phenotypes in MS.

## Results

### Generation of an iPSC collection for MS research

We generated 17 MS iPSC lines (6 RRMS, 6 SPMS, and 5 PPMS) from skin biopsies of people with MS^18^(Figure 1A). Participants with MS were categorized as as RRMS, PPMS or SPMS, according to the phenotypic classification system used at the time of enrollment (Figure 1B)^5^. Skin fibroblasts were reprogrammed by modified mRNAs using the NYSCF Global Stem Cell Array^®^ our fully automated system for generation of high-quality polyclonal iPSC lines that we have previously demonstrated to reduce technical variation^19^(Figure 1A, S1A). A total of 22 iPSC lines (including 5 healthy control iPSC lines) were used in this study. To characterize glial cells from healthy individuals and people with MS, we leveraged our previously published protocol^20,21^ to differentiate iPSCs into cultures enriched in astrocytes and oligodendrocyte lineage cells (encompassing all stages from oligodendrocyte progenitor cells to mature oligodendrocytes) (Figure 1C, Figure S1B-S1C).

**Figure 1.**
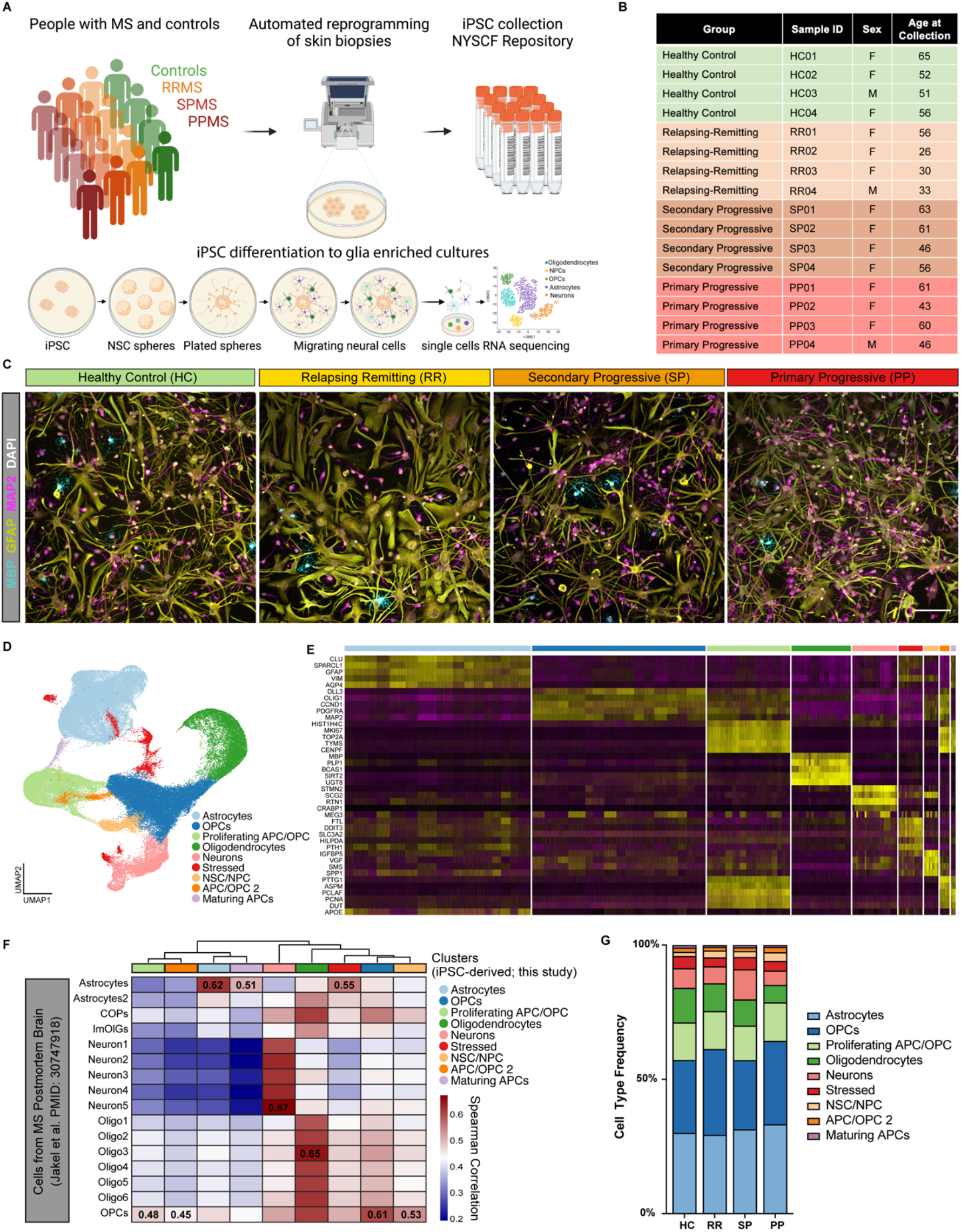
An iPSC-derived model to study CNS cell intrinsic dysfunction in MS. (A) Schematic representation of iPSC reprogramming from people with MS and healthy control skin biopsies and differentiation of iPSCs into glial CNS cultures. (B) Select demographic information for the iPSC lines used for scRNAseq analysis. (C) Representative images of iPSC-derived CNS cultures from healthy control and people with relapsing remitting MS, secondary progressive MS, and primary progressive MS. Cultures are stained for the mature oligodendrocyte marker MBP (teal), astrocyte marker GFAP (yellow), and neuron marker MAP2 (pink). Scale bar is 100µm. (D) UMAP of integrated single-cell analysis from 4 healthy control lines, 4 relapsing remitting lines, 4 secondary progressive lines, and 4 primary progressive lines, showing major cell type clusters. (E) Heatmap of the top 4 enriched genes for each cluster in (D). (F) Heatmap depicting the correlation between clusters in (D) and cell types from scRNAseq analysis of MS brain tissue (Jakel et al. PMID: 30747918). Spearman correlation values generated using the R package ClustifyR. (G) Distribution of cell types within iPSC-derived CNS cultures from healthy control (HC), relapsing remitting (RR), secondary progressive (SP), and primary progressive (PP) MS.

### Single-cell transcriptional profiling of iPSC-derived neural cells from people with MS

Using 16 iPSC lines (4 ctrl, 4 RRMS, 4 PPMS, 4 SPMS), we performed single-cell RNA sequencing on glia-enriched cultures, and we characterized a total of 122,228 cells with an average of 7639 ± 1141 cells per line (Figure S1D). Data were first filtered to remove doublets, low quality cells with less than 200 genes captured, and cells with >25% mitochondrial reads (Figure S1E-S1F). Unsupervised clustering of the remaining high-quality cells identified 9 cell clusters shared by all individual samples (Figure 1D, S1G). We next identified differentially expressed genes enriched in each cluster and used those to assign a cell type to each cluster (Table S1). The largest identified cell clusters were: astrocytes (*GFAP* and *AQP4*), oligodendrocyte progenitor cells (OPCs) (*OLIG1* and *PDGFRA*), proliferating progenitor cells (*MKI67* and *TOP2A*), oligodendrocytes (*MBP* and *PLP1*), and neurons (*STMN2* and SCG2) (Figure 1E, S1G). Our data were consistent with previously reported classifications of cell-specific clusters from human MS brain tissues^11,12^(Figure 1F, S1H). These data show that iPSCs from people with MS and healthy controls successfully differentiate into MS-relevant cell types and can be used to explore molecular and functional differences in MS that are intrinsic to glial cells.

### iPSC-derived glial enriched cultures from people with PPMS generate fewer oligodendrocytes

We next asked whether the frequency of cell types changed in iPSC-derived CNS cultures from people with MS compared to healthy controls. Surprisingly, we found a consistent impairment in the generation of oligodendrocytes in PPMS cultures (6.45%, 2061/31909) compared to healthy control cultures (12.86%, 4290/33344) (Figure 1G). To confirm that this was not caused by a general defect in the differentiation potential of the MS lines, we examined whether the generation of astrocytes and neurons was similarly affected. Using a sorting protocol for the CD49f^+^ astrocyte marker^22^, and by generating cortical neurons using a distinct protocol^23^, we did not detect any difference in the percentage of CD49f^+^ astrocytes or MAP2^+^ neurons generated from MS versus healthy control lines (Figure S1I-K).

To further explore the decreased oligodendrocyte formation in PPMS cultures we re-clustered only the iPSC-derived oligodendrocyte lineage cells and identified five populations representing different stages of differentiation of OPCs into oligodendrocytes (Figure 2A, Table S1). Three OPC clusters (34.31% OPCs.1, 27.19% OPCs.2, and 14.73% OPCs.3) expressing OPC markers, one cluster of newly formed oligodendrocytes (14.73%) expressing genes increased early in oligodendrocyte formation, and one cluster of mature oligodendrocytes (9.77%) expressing myelin genes (Figure 2A-D, Figure S2A). While all subtypes were present in all samples, the distribution of cell subtypes for each sample confirmed the decreased number of newly formed and mature oligodendrocytes, with a corresponding increase in OPCs only in PPMS cultures (Figure 2E-F).

**Figure 2.**
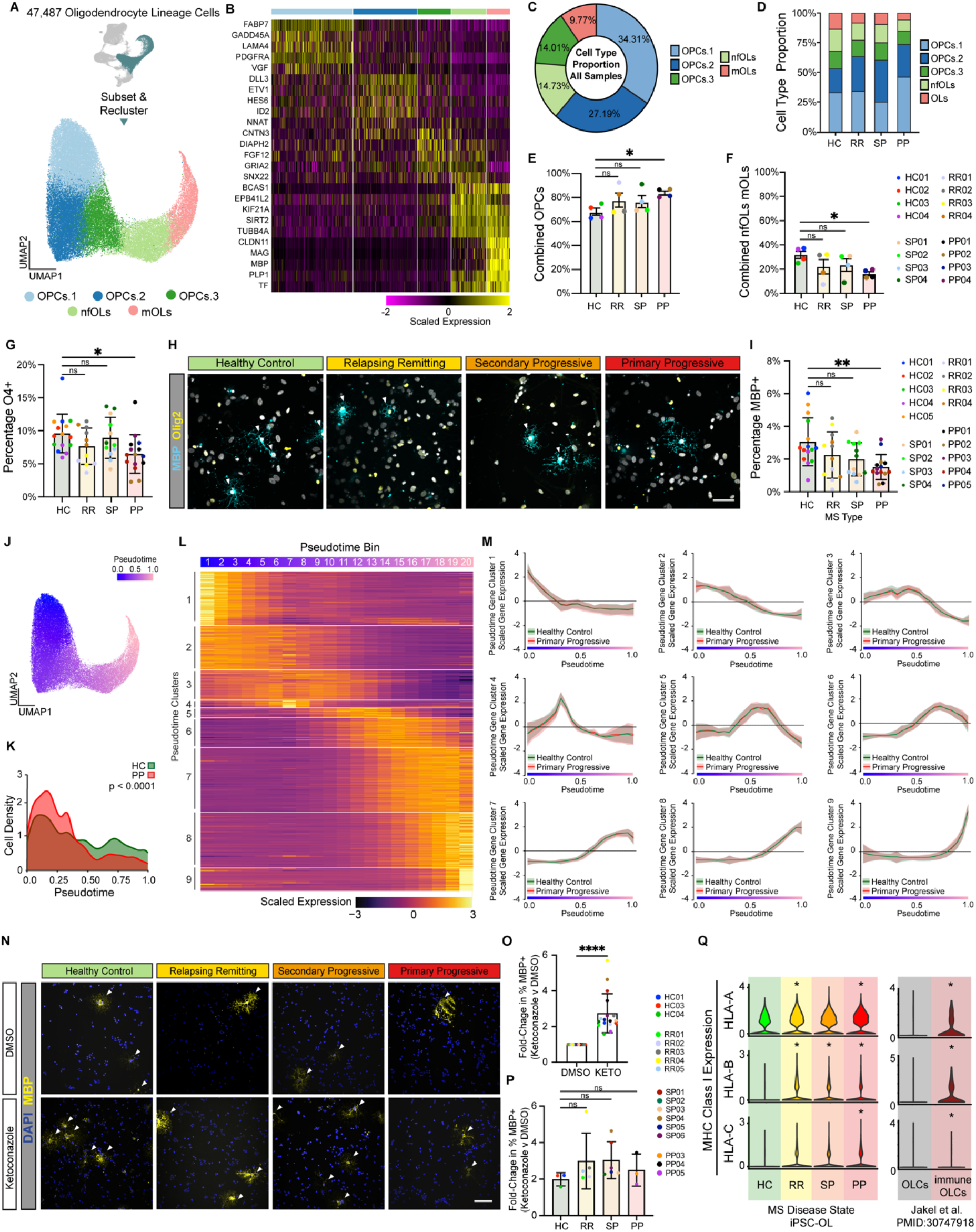
iPSC-derived cultures from people with MS generate fewer oligodendrocytes. (A) UMAP of 47,487 oligodendrocyte lineage cells subset and re-clustered. (B) Heatmap depicting the scaled expression of the top 5 enriched genes for each oligodendrocyte lineage cell cluster in Figure 2A. (C) The proportion of each oligodendrocyte lineage cell type in iPSC-derived cultures from all samples. (D) The proportion of each oligodendrocyte lineage cell type in iPSC-derived cultures from healthy control (HC), relapsing remitting (RR), secondary progressive (SP), and primary progressive lines. (E) The percentage of cells in iPSC-derived cultures from healthy control (HC), relapsing remitting (RR), secondary progressive (SP), and primary progressive lines that are oligodendrocyte progenitor cells (OPCs). The percentage of OPCs in PP cultures is significantly higher than HC cultures. Data presented as mean +/- s.e.m. for *n* = 4 per group. p-value generated by Welch’s ANOVA with Dunnett’s T3 correction for multiple comparisons against healthy controls (HC). (F) The percentage of cells in iPSC-derived cultures from healthy control (HC), relapsing remitting (RR), secondary progressive (SP), and primary progressive lines that are newly formed oligodendrocytes (nfOLs) or mature oligodendrocytes (mOLs). The percentage of combined nfOLs and mOLs in PP cultures is significantly lower than HC cultures. Data presented as mean +/- s.e.m. for *n* = 4 per group. p-value generated by Welch’s ANOVA with Dunnett’s T3 correction for multiple comparisons against healthy controls (HC). (G) The percentage of oligodendrocyte lineage cells (OLIG2^+^) in iPSC-derived cultures from healthy control (HC), relapsing remitting (RR), secondary progressive (SP), and primary progressive lines that are positive for the early oligodendrocyte marker O4. The percentage of OLIG2 positive oligodendrocyte lineage cells in PP cultures that are O4^+^ is significantly lower than in HC cultures. Error bars show mean ± standard deviation (n= 3 technical replicates per line for 4-5 lines per group). p-values generated by one-way Anova with Dunnett’s correction for multiple comparisons. (H) Representative images of iPSC-derived CNS cultures stained for the mature oligodendrocyte marker MBP (teal). Scale bar is 50µm. (I) The percentage of oligodendrocyte lineage cells (OLIG2^+^) in iPSC-derived cultures from healthy control (HC), relapsing remitting (RR), secondary progressive (SP), and primary progressive lines that are positive for the mature oligodendrocyte marker MBP. The percentage of OLIG2^+^ oligodendrocyte lineage cells in PP cultures that are MBP^+^ is significantly lower than in HC cultures. Error bars show mean ± standard deviation (n= 3 technical replicates per line for 4-5 lines per group). p-values generated by one-way ANOVA with Dunnett’s correction for multiple comparisons. (J) Pseudotime plot of oligodendrocyte lineage trajectory from OPCs to mature oligodendrocytes. (K) Cell density plot that shows the distribution of cells across the oligodendrocyte lineage trajectory from OPCs to mature oligodendrocytes for iPSC-derived cells from healthy controls (HC) or people with primary progressive MS (PP). p-value generated with a two-sample Kolmogorov-Smirnov test. (L) Heatmap depicting the scaled expression of genes that were determined to have pseudotime specific expression profiles. (M) Comparison of pseudotime gene expression profiles between healthy control and primary progressive oligodendrocyte lineage cells. (N) Expression of MHC Class I genes in healthy control (HC), relapsing remitting (RR), secondary progressive (SP), and primary progressive cultures from this study. Expression of MHC Class I genes in oligodendrocyte lineage cells (OLCs) and immunological OLCs from MS postmortem brain (Jakel et al. PMID: 30747918). p-value generated by Wilcoxon ranked sum test within the Seurat R package. * p < 0.05. (O) Representative images of healthy control (HC), relapsing remitting (RR), secondary progressive (SP), and primary progressive cultures treated with either vehicle (DMSO) or the oligodendrocyte enhancing compound ketoconazole at 100µm. (P) Fold-change in the percentage of MBP positive cells in all cultures treated with vehicle (DMSO) or ketoconazole (KETO) at 1µM for 24 hours. Error bars show mean ± standard deviation (n= 17 lines). p-values generated by two-way paired t-test. (Q) Fold-change in the percentage of MBP positive cells in healthy control (HC), relapsing remitting (RR), secondary progressive (SP), and primary progressive cultures treated with either vehicle (DMSO) or the oligodendrocyte enhancing compound ketoconazole. Data is presented as mean +/- standard deviation for *n* = 3-6 lines per group. p-values generated by one-way ANOVA with Dunnett’s correction for multiple comparisons.

Next, we performed orthogonal validation of oligodendrocyte formation using immunocytochemistry with antibodies specific for: the oligodendrocyte lineage cell marker OLIG2 (expressed at all stages in the oligodendrocyte lineage), O4, a marker of late adult progenitors and newly formed oligodendrocytes, and myelin basic protein (MBP) a major protein component of myelin and marker of mature oligodendrocytes. There was no difference between MS and control cultures in the percentage of total cells stained with OLIG2; however, in agreement with our scRNAseq data, PPMS cultures exhibited a significant decrease in cells stained for either the immature oligodendrocyte marker O4 or the mature oligodendrocyte marker MBP (Figure 2G-I, S2B). We then performed pseudotime analysis to place oligodendrocyte lineage cells along the differentiation trajectory from OPCs to mature oligodendrocytes (Figure 2J). Density analysis again confirmed that PPMS cultures contained fewer oligodendrocytes, but more OPCs (Figure 2K). We next generated gene modules with distinct expression patterns along the differentiation trajectory of OPCs to oligodendrocytes (Figure 2L, Table S2). Despite PPMS cells showing a delay in the progression toward mature oligodendrocytes, we could not detect differences in the expression of these oligodendrocyte differentiation gene modules (Figure 2M). Furthermore, the size of arborized mature oligodendrocytes was not reduced in PPMS cultures (Figure S2D,E). Together these data suggest that PPMS iPSC lines generate fewer mature oligodendrocytes, but morphology and gene expression of the cells that do form are not altered.

### iPSC-oligodendrocytes respond to myelinating drugs independently of disease state

Promoting mature oligodendrocyte formation and remyelination are major clinical goals in MS^24^. To this end, multiple groups have identified small molecule enhancers of oligodendrocyte formation^25–28^. Nevertheless, it is not known whether OPCs from people with MS are responsive to oligodendrocyte-enhancing therapies, or whether OPCs from distinct MS types might respond differently to drugs that promote oligodendrocyte formation. To test this, we treated iPSC-derived cultures from healthy control and MS individuals with ketoconazole and TASIN-1, two small molecules that we have previously identified promote oligodendrocyte formation by inhibiting specific enzymes in the cholesterol biosynthesis pathway^29^. We found that treatment with either ketoconazole or TASIN-1 increased the formation of MBP^+^ mature oligodendrocytes in all MS cultures (Figure 2N-P, S2G), showing that MS cells are responsive to potential remyelinating therapies regardless of MS type.

### iPSC-derived MS oligodendrocytes exhibit an immune-like transcriptome profile

Recent reports have described immunological oligodendrocyte lineage cells in mouse models of MS and in human MS tissue that may contribute to oligodendrocyte lineage cell cytotoxicity, decreased oligodendrocyte formation, and ultimately failed remyelination^11,30,31^. Even in the absence of inflammatory stimuli or activation by peripheral immune cells, we found that iPSC-derived oligodendrocyte lineage cells from people with MS expressed increased levels of MHC class I transcripts, with oligodendrocyte lineage cells from PPMS expressing significantly higher levels of all three HLA-A, HLA-B, and HLA-C (Figure 2Q, S2F). This mirrors the increased expression of MHC class I transcripts found in immunological oligodendrocyte lineage cells from single-nuclei RNAseq analysis of MS brain tissue and may contribute to the decreased formation of oligodendrocytes in iPSC-derived CNS cultures from people with PPMS.

### iPSC-derived MS astrocytes exhibit an immune and inflammatory profile

Mounting evidence shows that reactive astrocytes contribute to the development and progression of MS pathology^32^. To explore potential phenotypes specific to MS astrocytes, we next subselected and re-clustered astrocytes from iPSC-derived healthy control and MS cultures (Figure 3A-B, S3A, Table S1). We identified eight distinct subtypes of astrocytes in iPSC-derived CNS cultures, with astrocyte cluster 6 consisting almost exclusively of cells from individuals with MS (Figure 3C-D, S3B). Gene Ontology analysis of astrocyte cluster 6 defining genes showed enrichment for antigen processing and presentation, inflammatory signaling, and Epstein-Barr virus (EBV) infection (Figure 3E). In addition, astrocyte cluster 7 was enriched with cells from RRMS and SPMS cultures (Figure S3C) and genes defining astrocyte cluster 7 were associated with inflammation including cytokine, interferon, and NFkB signaling terms (Figure S3D). We also found that MS cultures were depleted for cells in astrocyte cluster 3 (Figure S3E). Gene Ontology analysis of genes defining the astrocyte cluster 3 showed enrichment for axon development, neuron development, and sterol catabolism (Figure S3F). To validate our single-cell RNAseq results we used RNAscope *in situ* hybridization to localize MS-enriched gene expression in culture. This approach confirmed that HLA class II histocompatibility antigen, DR alpha chain (*HLA-DRA*, involved in antigen presentation) is increased in PPMS cultures compared to healthy controls (Figure 3F-H). Together these data suggest that, even in the absence of any inflammatory stimuli, astrocytes from people with MS are more likely to acquire an inflammatory state, at the expense of a neuro-supportive state, and that this may contribute to MS disease progression.

**Figure 3.**
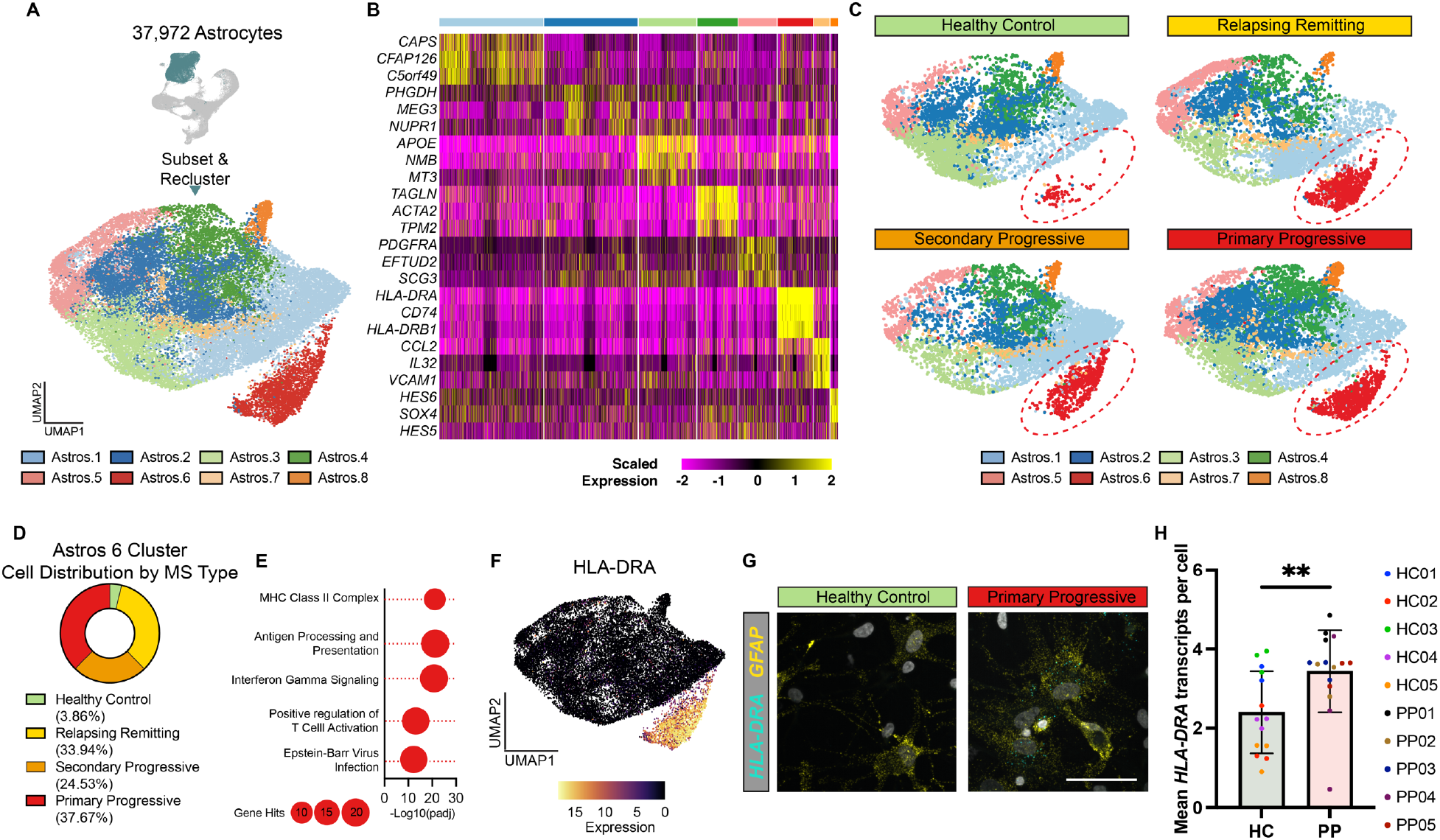
scRNAseq reveals a reactive astrocyte subtype enriched in iPSC-derived MS cultures. (A) UMAP of 37,927 astrocytes subsetted and reclustered. Eight unique astrocyte subclusters were identified. (B) Heatmap showing the scaled expression of the top three genes enriched in each astrocyte subcluster. (C) UMAP plots of healthy control, relapsing remitting, secondary progressive, and primary progressive iPSC-derived astrocytes. The Astro.6 cluster is enriched only in iPSC-derived astrocytes from patients with MS and not healthy controls. (D) Distribution of healthy control, relapsing remitting, secondary progressive, and primary progressive iPSC-derived astrocytes in the astrocyte subcluster 6. (E) Gene ontology analysis of genes significantly increased in astrocytes subcluster 6 compared to all other astrocyte subclusters. (F) UMAP plot overlayed with the expression of *HLA-DRA*, a gene significantly increased in Astro.6 compared to all other astrocyte clusters. (G) Representative images of RNAscope *in situ* hybridization for *GFAP* and *HLA-DRA* in iPSC-derived cultures from healthy control and primary progressive patients. Images show localization of *HLA-DRA* to *GFAP^+^* astrocytes in primary progressive cultures only. Scale bar, 50μm. (H) Quantification of RNAscope *in situ* hybridization for *GFAP* and *HLA-DRA* in iPSC-derived cultures from healthy control (HC) and primary progressive (PP) patients. Error bars show mean ± standard deviation (n= 2-3 technical replicates per line for 5 lines per group). p-value generated by one-way unpaired t-test.

### iPSC-derived MS astrocytes mirror pathological astrocytes in MS brains

It has been shown that astrocytes can adopt an inflammatory pathological state in response to the microglial derived cytokines TNF, IL1α, and C1q (TIC)^22,33,34^. These TIC-induced neurotoxic reactive astrocytes have been observed in most neurodegenerative diseases including MS, Alzheimer’s, Parkinson’s, and amyotrophic lateral sclerosis^33,35^. Analysis of human iPSC-astrocytes has also shown that cells containing a common MS risk SNP express higher levels of genes associated with TIC-induced astrocyte reactivity^36^. We therefore sought to investigate whether, even in the absence of inflammatory stimuli, the iPSC-derived astrocytes in MS cultures reflect a similar pathological state. To do this, we integrated iPSC-astrocytes from healthy controls and people with MS, with single-cell analysis of healthy control human iPSC-astrocytes treated with TIC cytokines^22,23^. Unbiased clustering of this integrated data set identified 12 distinct astrocyte clusters (Figure S4A, Table S3), three of which (clusters 8, 9, and 10), almost exclusively contained iPSC-derived astrocytes from people with all clinical subtypes of MS, generated in this study, and TIC-treated iPSC-derived astrocytes from healthy individuals (Figure S4B-C, S4D, S4F, S4H). Moreover, Gene Ontology analysis of genes defining these clusters showed enrichment for terms associated with inflammation and immune processes including antigen processing and presentation, cytokine signaling, and again EBV infection (Figure S4E, S4G, S4I). Together these data show that iPSC-derived astrocytes from people with all clinical subtypes of MS transition to a pathological reactive state, independent of exogenous inflammatory stimuli or interaction with immune cells.

Finally, we sought to determine whether the iPSC-derived astrocyte subtype enriched in cultures from people with MS represents a pathological astrocyte state found in the MS brain. To do this, we integrated iPSC-derived astrocytes from healthy and MS cultures with publicly available single-nucleus analysis of astrocytes in MS brain tissues^12^. Integration of our iPSC-derived cultures with MS brain tissue data resulted in the identification of 7 distinct integrated astrocyte clusters (Figure 4A, Table S3). Two clusters, integrated astrocytes clusters 4 and 5 contained almost exclusively cells from MS brains and MS iPSC-derived cultures (Figure 4B-C). We focused on integrated astrocyte cluster 5 that contained cells from all clinical subtypes of MS, while cluster 4 was driven by cells from only two samples. Differential gene expression analysis identified MHC Class I and Class II genes as significantly enriched in integrated astrocyte cluster 5 compared to other clusters. Moreover, Gene Ontology analysis of genes specific to integrated astrocyte cluster 5 showed enrichment for antigen processing and presentation, inflammatory signaling, neurodegeneration, and again EBV infection terms (Figure 4E-F). These data show that even in the absence of inflammatory stimuli or peripheral immune activation, iPSC-derived CNS cultures from people with MS generate pathological astrocyte subtypes that mirror those identified in MS brains.

**Figure 4.**
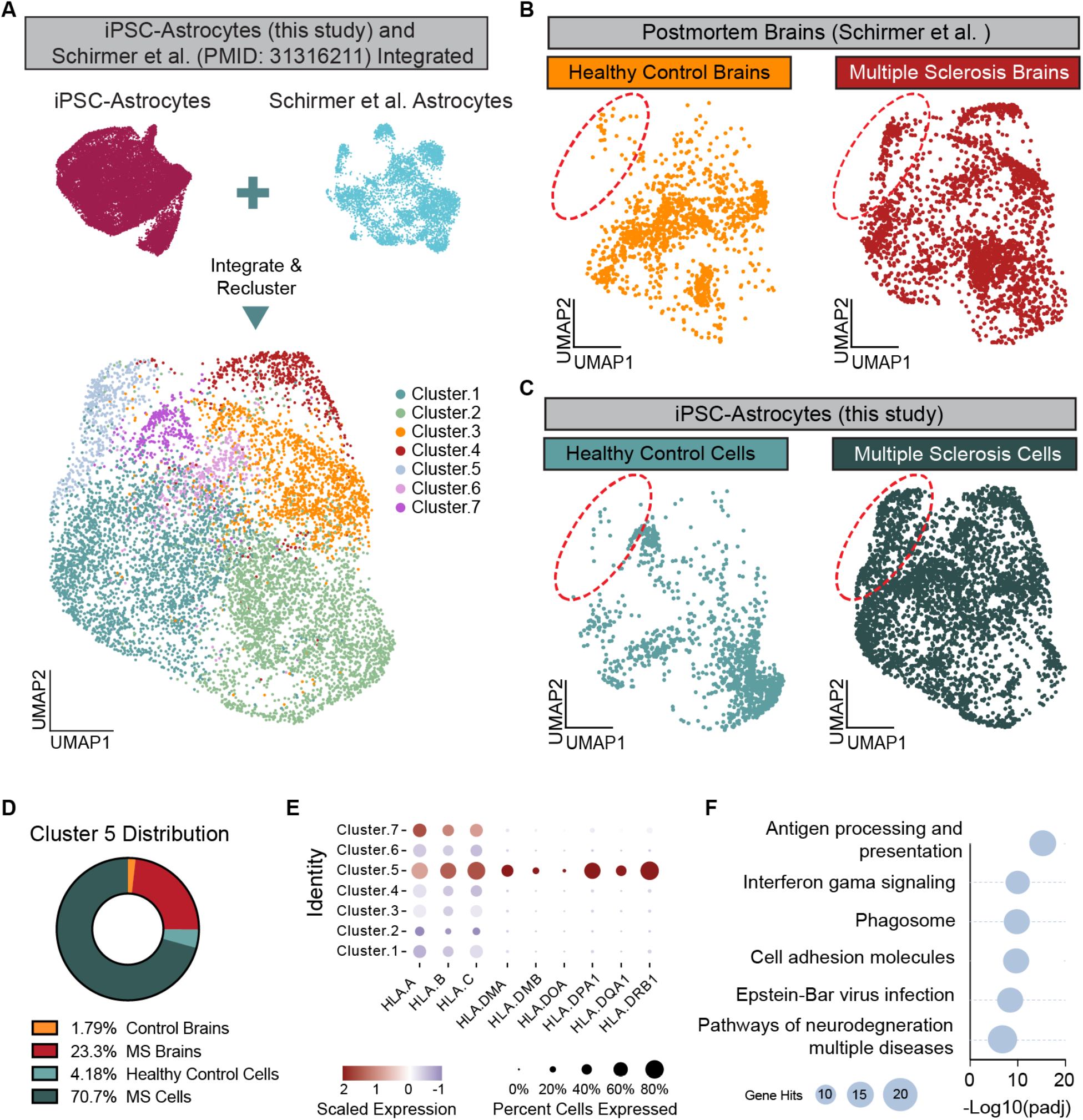
iPSC-derived astrocytes from people with MS mirror astrocytes from MS brains. (A) UMAP plot of astrocytes from this study integrated with astrocytes from healthy control individuals and people with MS (Schirmer et al. PMID: 31316211). (B) UMAP plots showing the distribution of cells from healthy control brains and MS brains. Red circle highlights Cluster 5 which is enriched for cells from MS brains. (C) UMAP plots showing the distribution of cells from iPSC-derived astrocytes from healthy control individuals and individuals with MS. Red circle highlights Cluster 5 which is enriched for iPSC-derived astrocytes from people with MS. (D) Distribution of cells from healthy control brains (yellow), MS brains (red), healthy control iPSC-derived astrocytes (light blue), and MS iPSC-derived astrocytes (dark blue) in Cluster 5 of the integrated data sets. (E) Dot plot showing the scaled expression of some MHC Class I and Class II genes in each of the clusters from the integrated data sets. (F) Gene ontology analysis of the top 100 genes enriched in Cluster 5 of the integrated data set compared to all other clusters in the integrated data set.

## Discussion

MS is a complex disease resulting from the interaction between environmental factors, for example EBV infection, and genetic predisposition^3^ associated with peripheral immune and glial cells^2,14^. The role of glial cell-intrinsic dysfunction in the initiation and progression of MS remains unclear^37,38^. Here we leveraged the unique opportunity iPSC technologies offer to address this issue by investigating human glial cells independent of the complex *in vivo* environment, which is chronically altered by inflammation and infiltration of peripheral immune cells.

A handful of studies have investigated iPSC-derived neural progenitors and oligodendrocytes from people with MS; these studies used a limited number of lines (up to 3) and have produced conflicting reports on MS phenotypes in CNS cells^39–43^. Leveraging the NYSCF automated platform for iPSC derivation, we generated 22 lines including healthy controls, RRMS, SPMS and PPMS samples, differentiate them into glia-enriched cultures and performed single-cell transcriptome analysis. Notably, single cell transcriptome profiles revealed intrinsic features of MS oligodendrocytes and astrocytes, including decreased numbers of oligodendrocytes in PPMS cultures. Mature oligodendrocytes in our cultures do not commit to myelination and therefore undergo cell death, similarly to what occurs *in vivo*^44,45^. Therefore, we cannot exclude that the lower number of oligodendrocytes is a consequence of increased cell death. This hypothesis is substantiated by emerging findings implicating structural myelin abnormalities as initial triggers of inflammation and demyelination^46–48^, pointing to putative degenerative processes intrinsic to oligodendrocytes. This hypothesis will be tested in the future, as soon as platforms to efficiently reproduce human myelination *in vitro* become available. We also found increased immune and inflammatory gene expression in oligodendrocyte lineage cells and astrocytes that mirrors transcriptional profiles of these glia in post-mortem MS brains. The transition of oligodendrocyte lineage cells to an immune-like state has been described in postmortem MS brains^11,30,31^; nevertheless the etiology of these cells and their effect on remyelination in MS remains unclear. Given that iPSC-derived oligodendrocyte lineage cells in our cultures were never exposed to inflammation or peripheral immune cells, our findings indicate that in MS, oligodendrocyte lineage cells are intrinsically primed to transition to an immune-like state.

In conclusion, our study demonstrates that iPSC-derived glial enriched cultures from people with MS are a powerful model to identify CNS-intrinsic phenotypes in MS. Moving forward, future studies of iPSC-derived glia from people with MS could help identify disease mechanisms that explain, for example, why some people present with elevated inflammation and multiple relapses, while others develop a progressive course with minimal inflammatory activity. Importantly, these findings could reveal novel glia-specific targets for urgently needed therapeutics that stop or reverse disease progression in MS.

## Limitations

While this study includes the largest cohort of iPSCs from people with MS reported to date it is still too small to identify genotype-phenotype associations. The differentiation protocol used in these studies does not generate cultures with compact myelin wrapping axonal fibers or with microglia. Additional studies with myelinated cultures and iPSC-derived microglia from people with MS are warranted.

## Data and Resource Availability

All sequencing datasets generated in this study have been deposited in Gene Expression Omnibus (https://www.ncbi.nlm.nih.gov/geo/) under accession code GSE238221.

Reviewer access code: ubwjseoivdcbjkx

We have genereated a collection of MS iPSC lines to enable investigation of molecular mechanisms of CNS dysfunction and potential glial targets for therapeutic intervention. NYSCF Research Institute iPSC lines may be made available on request through the NYSCF Research Institute repository (nyscf.org/repository).

## Acknowledgments

This study was supported by grants from the Department of Defense Congressionally Directed Medical Research Programs (CDMRP) Multiple Sclerosis Research Program (MSRP), Award number DOD W81xWH-15–1–0448 (P.C.); the Corinne Goldsmith Dickinson Center for Multiple Sclerosis (V.F.); the New York State Stem Cell Science (NYSTEM), Award number C32586GG (V.F.); the National Multiple Sclerosis Society Career Transition Award TA-2105-37619 (B.L.L.C); the National Institutes of Health, R35NS116842 (P.J.T.) and R35NS111604 (P.C.), and sTF5 Care (P.J.T.); and by the New York Stem Cell Foundation Research Institute. We thank Dr. Raeka Aiyar for manuscript editing. We are extremely grateful to the people that donated the skin biopsies for this study.

## Author Contributions

Conceptualization, B.L.L.C., L.B, P.J.T., and V.F.; Methodology, B.L.L.C., L.B; Validation, B.L.L.C., L.B; Formal Analysis, B.L.L.C., L.B; Investigation, B.L.L.C., L.B, M.S., T.R., B.M.; K.K.; D.M.; D.P.; K.B.; Resources, P.C, I.K.S, P.J.T., V.F., Data Curation, B.L.L.C.; Writing-original draft, B.L.L.C., L.B., P.J.T., and V.F.; Writing-Review & Editing, all co-authors.; Project Administration, P.J.T., V.F., Funding Acquisition, P.J.T., P.C., V.F.

## METHODS

### iPSC generation, quality control, and culture conditions

iPSC lines were derived by reprogramming fibroblasts from skin biopsies. Dr. Ilana Katz Sand at the Corinne Goldsmith Dickinson Center for MS at Mount Sinai recruited MS patients, and performed clinical characterization to distinguish RRMS, PPMS and SPMS forms (registered clinical trial NCT02549703). The original study also involved collection of cerebrospinal fluid^18^, and other MS-specific assignments and thus excluded healthy individuals. Control iPSC lines for were chosen from the NYSCF Research Institute repository, selecting for age- and sex-matched individuals (registered clinical trial NCT04270604). All iPSC lines were reprogrammed via modified mRNA technology, using the NYSCF Global Stem Cell Array®, a fully automated reprogramming process that minimizes line-to-line variability^19^. iPSCs were expanded onto Matrigel-coated dishes in mTeSR1 medium (StemCell Technologies) and passaged using enzymatic digestion with Stempro Accutase (ThermoFisher; A1110501) for 3-5 minutes and re-plated in mTeSR1 medium with the addition of 10μM ROCK Inhibitor (Y27632, Stemgent) for the first 24 hours. A total of 22 iPSC lines were used in this study. For each line a certificate of analysis (CoA) is provided, which includes the results for the following tests: sterility check, mycoplasma testing, karyotyping, identity test, and pluripotency check. Table 1 summarizes the demographic information of the individuals that donated the skin biopsies. Additional clinical information and whole genome sequencing of the MS lines may be provided on request.

**Table 1:** List of iPSC lines used in this study.

| Code | Repository Line name | Age at time of biopsy | Sex | Ethnicity | Disease state |
| --- | --- | --- | --- | --- | --- |
| HC01 | 050659-01-MR | 65 | F | Ashkenazi | Healthy control |
| HC02 | 051121-01-MR | 52 | F | Caucasian | Healthy control |
| HC03 | 050743-01-MR | 51 | M | Caucasian | Healthy control |
| HC04 | 051104-01-MR | 56 | F | Caucasian | Healthy control |
| HC05 | 051106-01-MR | 57 | F | Caucasian | Healthy control |
| RR01 | AK0007-01-MR | 56 | F | African-American | Relapsing remitting MS |
| RR02 | AK0028-01-MR | 26 | F | Caucasian | Relapsing remitting MS |
| RR03 | AK0024-01-MR | 30 | F | Caucasian | Relapsing remitting MS |
| RR04 | AK0014-01-MR | 33 | M | Caucasian | Relapsing remitting MS |
| RR05 | AK0013-01-MR | 54 | M | Caucasian | Relapsing remitting MS |
| RR06 | AK0027-01-MR | 69 | F | Caucasian | Relapsing remitting MS |
| SP01 | AK0015-01-MR | 63 | F | Caucasian | Secondary progressive MS |
| SP02 | AK0026-01-MR | 61 | F | Caucasian | Secondary progressive MS |
| SP03 | AK0008-01-MR | 46 | F | Caucasian | Secondary progressive MS |
| SP04 | AK0011-01-MR | 56 | F | Caucasian | Secondary progressive MS |
| SP05 | AK0005-01-MR | 65 | F | Caucasian | Secondary progressive MS |
| SP06 | AK0012-01-MR | 51 | F | African-American | Secondary progressive MS |
| PP01 | AK0001-01-MR | 61 | F | Caucasian | Primary progressive MS |
| PP02 | AK0009-01-MR | 43 | F | Caucasian | Primary progressive MS |
| PP03 | AK0004-01-MR | 60 | F | Caucasian | Primary progressive MS |
| PP04 | AK0010-01-MR | 46 | M | Caucasian | Primary progressive MS |
| PP05 | AK0003-01-MR | 52 | F | Caucasian | Primary progressive MS |

### iPSC differentiation into glia-enriched cultures

Cells were grown at 5% CO_2_ in a 37°C incubator. hiPSCs were induced along the neural lineage and differentiated using our previously published protocol^21^. Briefly, hiPSCs were plated at 1-2 × 10^5^ cells per well on matrigel-coated six-well plates in mTeSR1 medium with 10 μM ROCK inhibitor Y27632 (Stemgent; 04-0012). Starting the following day, cells were fed daily with mTeSR1 medium (StemCell Technologies; 85850). Once distinct colonies of ~200 µm in diameter were formed, differentiation was induced (day 0). Details for the differentiation protocol can be found in our previous publications^22,49^ and a schematic with media composition is provided in figure S1. Around d30 of differentiation, OLIG2^+^ neurospheres were plated into polyOrnithine/laminin-coated dishes to allow cell attachment and migration. Radial glial cells anchored the sphere to the well and progenitor migrated out, generating a neurons, astrocytes and oligodendrocytes in this precise order. At the end of the differentiation, between day 70 and day 85, cultures were dissociated using different protocols depending on downstream analyses. For exposure to myelinating compounds, RNAscope and immunofluorescence analyses, cells were incubated for ~25 minutes with StemPro Accutase (ThermoFisher; A1110501) and passed through a 70µm strainer. The resulting single-cell suspension was re-plated or sorted for CD49f (BD Biosciences; 555736) to purify CD49f positive astrocytes and enrich oligodendrocyte lineage cells within the CD49f negative fraction. The FACS protocol has been described in details previously^23^. Single cell suspensions were re-plated onto PO/Lam coated plates and 24 hours after plating, medium was switched to “glial medium”, and cells were fed with two-third media changes every other day. For scRNAseq analyses, cultures were dissociated using papain, as detailed below.

### iPSC differentiation into cortical neurons

hiPSCs were plated in a 12-well plate in mTeSR1 media with 10μM ROCK inhibitor Y2732. Starting the next day (d0), cells were fed daily with neural induction media with SB431542 (20 μM; Stemgent, 040010), LDN193189 (100nM; Stemgent, 040074), XAV939 (1 μM; Tocris, 3748). Neural induction media consisted of 1:1 DMEM/F12 (ThermoFisher; 11320033) and Neurobasal (ThermoFisher; 21103049) with 1x Glutamax (Thermo Scientific 35050061), 1x N2 supplement (Gibco 17502-048), and 1x B27 supplement without Vitamin A (Gibco; 12587-010). On day 10, the media was switched to neural induction media with XAV939 (1μM), with continuing daily media changes. On day 15, cells were dissociated using Accutase and either frozen in Synth-a-freeze, or plated in neuronal media at 50k/well in a PO/Lam coated 96-well plate (Corning; 353376). Neuronal medium consist of Neurobasal (StemCell Technologies; 05790) with 1xB27 supplement (ThermoFisher; 17504001), and 10 μM ROCK inhibitor. On day 16, the media was switched to neuronal media with BDNF (40ng/mL; R&D Systems, 248-BDB), GDNF (40ng/mL; R&D Systems 212-GD), Laminin (1μg/mL), dbcAMP (250 μM), ascorbic acid (200 μM), PD0325901 (10 μM; Reprocell, 04-0006), SU5402 (10 μM; Sigma-Adrich SML0443), DAPT (10 μM; Tocris, 2634), and ROCK inhibitor (10μM)^50^. Cells were fed every other day. Starting on day 25, cells were fed every other day with neuronal medium with BDNF (40ng/mL), GDNF (40ng/mL), Laminin (1mg/mL), dbcAMP (250 μM), ascorbic acid (200 μM). This differentiation protocol has been thoroughly described in our previously published protocol^23^, with the following changes: here Neurobasal was used instead of Brainphys after day 15, and PD0325901, SU5402, DAPT, and ROCK inhibitor were taken out of the media at day 25.

### Ketoconazole and TASIN-1 treatment, immunostaining and analysis

CD49f negative cells (enriched of oligodendrocyte lineage cells) were seeded at 25K cells per well (2-3 replicates minimum) onto Poly-ornithine/laminin-coated 96 well Perkin Elmer Cell Carrier Ultra (CCU) microplates at day 75-80. Cells were cultured for 7 days in glia maturation media and treated with either vehicle (DMSO, 0.1%V/V), Ketoconazole (1μM) or TASIN-1 (0.1uM, kindly provided by Drew Adams, Case Western Reserve University) with feeds every other day, a total of 3 treatments. On Day 8, cells were fixed in 2% PFA/PBS solution for 10 minutes. Permeabilization and blocking were done in PBS with 0.1% Saponin and 5% Normal Donkey Serum or Normal Goat Serum. For the ketoconazole experiment, cells were labeled with MBP. For the TASIN-1 experiment, we included SOX10 and Olig2. All image acquisition was done with the Opera Phenix High Content Screening system (Perkin Elmer) in confocal-mode. For the 96-well CCU plates, we collected 25 images per well at 10X magnification which included an average of 9,000 cells per well. NUNC plates are not compatible with the Phenix and were modified with a 3D printed part. Using the Opera Phenix, we acquired 60 fields, 4 Z-planes covering 21 µm at 10X magnification per well which included an average of 50,000 cells around each sphere. All analysis was done using the Harmony software (Perkin Elmer). In summary, the analysis first traced intact nuclei based on the DNA stain fluorescence and then selected nuclei that were larger than 40-50μm^2^ surface area and had intensity levels lower than the brightness of pyknotic nuclei. We scored the MBP positive cells by identifying the cell type marker positive surrounding region around the nuclei and selected cells based on the mean fluorescence intensity in the surrounding ROI. Because the CD49f negative fraction is highly heterogeneous, we included a step in the analysis that clustered nuclei by distance to minimize object splitting and overcounting of MBP^+^ oligodendrocytes. For the TASIN-1 experiments, we used nuclear SOX10 and Olig2 labels to further filter oligodendrocyte lineage cells before identifying MBP oligodendrocytes. Percentage was calculated at well level: Total MBP positive cells/Total nuclei X 100 in well. Drug treated well replicates were averaged for the fold change ratio calculations. Plots were generated with Prism analysis software.

### Cell dissociation and library preparation for single-cell RNA sequencing (scRNAseq)

Day 85 cultures from 16 lines were detached enzymatically in parallel using papain (Worthington; LK003153) and were filtered through 40µm Flowmi Cell Strainers (Scienceware; H13680-0040) to obtain a single cell suspension. Single cells were processed using the 10X Single Cell 30 v3.1Rev Bprotocol. Briefly, we loaded the Chromium Single Cell Chip G (10X Genomics; PN-2000177) with 7,000 cells/sample and we performed library preparation as per the Chromium Single Cell 30 Library & Gel Bead Kit manufacturer’s recommendations (10X Genomics; PN-120237 and PN1000121). We used the Chromium i7 Multiplex Kit (10X Genomics; PN-120262). Quality control was performed using the Qubit 4 Fluorometer (ThermoFisher; Q33227) and the Agilent 4200 TapeStation system. The resulting cDNA library was sequenced on a NovaSeq/HiSeq 2×150 bp, and 50,000 reads per cell were obtained.

### scRNAseq data processing

Sequence data were first processed by 10x Cell Ranger v3.0.2 to align reads to the human transcriptome build GRCh38, remove empty droplets, remove droplets containing multiple cells, and generate a feature-barcode matrix. Preprocessing was then performed with Seurat v3.2^51^. Following the standard pre-processing tutorial for the Seurat analysis package (https://satijalab.org/seurat/archive/v2.4/pbmc3k_tutorial.html). For each individual sample cells with fewer than 200 genes and/or percent mitochondrial reads above 25% were removed.

### Identification of broad shared cell types across integrated scRNAseq samples

In order to first identify broad cell types that were shared across all of the samples we performed integration of the data at the sample level. Integration was performed with Seruat v3.2^52^ following the standard integration tutorial (https://satijalab.org/seurat/articles/pbmc3k_tutorial.html). Each individual sample was log-normalized and variable features were identified using “NormalizeData” and “FindVariableFeatures” respectively. Integration anchors were then identified using “FindIntegrationAnchors” afterwhich the individual samples were integrated using “IntegrateData”. The integrated data set was then scaled using “ScaleData” while using difference in cell cycle score, RNA count, and percent mitochondrial reads as variable to regress.

Differential gene expression analysis was performed in Seurat v3.2 using “FindMarkers” with the Wilcoxon ranked sum test to identify differentially expressed genes between broad cell types from RRMS v Health Control, SPMS v Healthy Controls, and PPMS v Healthy Controls.

### scRNAseq analysis of oligodendrocytes

Analysis of oligodendrocyte lineage cells was performed in Seurat v3.2. Oligodendrocyte lineage cells were subsetted from the full data. These cells were then reanalyzed without integration to identify differences between samples and disease conditions. Data was log-normalized using “NormalizeData” and the top 2000 variable features were identified using “FindVariableFeatures”. The data set was then scaled using “ScaleData” while using difference in cell cycle score, RNA count, percent mitochondrial reads, and culture derivation batch as variable to regress. Principal component analysis was then ran using “RunPCA” with npcs = 50, followed by clustering using “FindNeighbors” dims = 1:11, and “FindClusters” with resolution = 0.4. Finally, “RunUMAP” was performed with dims = 1:11 to generate UMAP plots.

Differential gene expression analysis was performed in Seurat v3.2 using “FindMarkers” with the Wilcoxon ranked sum test to identify differentially expressed genes between oligodendrocyte lineage cells from RRMS v Health Control, SPMS v Healthy Controls, and PPMS v Healthy Controls.

### scRNAseq psuedotime analysis

Pseudotime analysis was performed using the Bioconductor package SCORPIUS (https://github.com/rcannood/SCORPIUS)^53^ which in comparison of multiple single-cell trajectory inference methods was identified as one a few methods that performed well under all conditions^54^. Seurat UMAP embeddings for the oligodendrocyte lineage cells was used to infer a trajectory by calling “infer_trajectory”. Candidate pseudotime genes were then called by using the Random Forest algorithm to rank genes according to their ability to predict the inferred trajectory of cells. Cells were then separated into 20 pseudotime bins across the oligodendrocyte lineage and the average scaled expression of candidate pseudotime genes for cells in each bin was calculated using the Seurat command “AverageExpression” and the result plotted on a heatmap ordered by pseudotime to identify pseudotime expression modules. This was done for both healthy control cells and cells from people with primary progressive MS to identify any changes in the expression pattern of pseudotime genes.

### scRNAseq analysis of astrocytes

Analysis of astrocytes was performed in Seurat v3.2. Astrocytes were subsetted from the full data and analyzed with integration across culture derivation batches and using SCTransform normalization (https://satijalab.org/seurat/articles/integration_introduction.html-performing-integration-on-datasets-normalized-with-sctransform-1)^55^ because exploratory analysis showed significant batch effects. Following SCTransform normalization data was integrated across batches by using “SelectIntegrationFeatures” to select 1000 integration features, integration anchors were found with “FindIntegrationAnchors” with k.anchor = 20, and finally “IntegrateData” was ran with normalization.method = “SCT”. Principal component analysis was then performed on the integrated data using “RunPCA” and npcs = 50, followed by clustering using “FindNeighbors” dims = 1:19, and “FindClusters” with resolution = 0.3. Finally, “RunUMAP” was performed with dims = 1:19 to generate UMAP plots.

Differential gene expression analysis was performed in Seurat v3.2. First “PrepSCTFindMarkers” was ran to prepare the SCTransform normalized data set for differential gene expression analysis. Then “FindMarkers” with the Wilcoxon ranked sum test was used to identify genes that are significantly enriched in each cluster compared to the data set as a whole.

### scRNAseq Integration of astrocytes with public data

Single-cell astrocyte data generated in this study were integrated with available single-cell raw read counts and metadata for single-cell RNA of iPSC-derived astrocytes from healthy controls exposed to TNF, IL1a, and C1q^22^. Integration was performed using SCTransform integration in Seurat. Briefly, both data sets were downsampled to 1500 astrocytes per group to ensure differences in cell number between groups didn’t affect integration. SCTransform normalization was then performed, after which data was integrated across batches by using “SelectIntegrationFeatures” to select 3000 integration features, integration anchors were found with “FindIntegrationAnchors” with k.anchor = 5, and finally “IntegrateData” was ran with normalization.method = “SCT”. Principal component analysis was then performed on the integrated data using “RunPCA” and npcs = 50, followed by clustering using “FindNeighbors” dims = 1:25, and “FindClusters” with resolution = 0.5. Finally, “RunUMAP” was ran with dims = 1:25 to generate UMAP plots.

Single-cell astrocyte data generated in this study were also integrated with available single-cell data of astrocytes from post-mortem brain tissue from people with MS^12^. The raw read counts and metadata for single-cell RNAseq from post-mortem MS brain tissue was acquired from the UCSC cell browser^56^. Single-cell astrocyte data from this study were integrated with MS astrocytes using SCTransform integration in Seurat. Briefly, both data sets were downsampled to 1250 astrocytes per group to ensure differences in cell number between groups didn’t affect integration. SCTransform normalization was then performed, after which data was integrated across batches by using “SelectIntegrationFeatures” to select 3000 integration features, integration anchors were found with “FindIntegrationAnchors” with k.anchor = 5, and finally “IntegrateData” was ran with normalization.method = “SCT”. Principal component analysis was then performed on the integrated data using “RunPCA” and npcs = 50, followed by clustering using “FindNeighbors” dims = 1:25, and “FindClusters” with resolution = 0.2. Finally, “RunUMAP” was ran with dims = 1:25 to generate UMAP plots.

For both integrated data sets, differential gene expression analysis was performed in Seurat v3.2 using “FindMarkers” with the Wilcoxon ranked sum test to identify differentially expressed genes between unbiased clusters.

### Immunofluorescence analysis

At the end of the differentiation, unsorted cells were plated onto 96-well plates to perform immunofluorescence analysis. Cells were fixed in 4% PFA for 10 minutes, washed 3x in PBS, and stored at 4°C. For staining, cells were incubated for one hour at room temperature in blocking solution consisting of PBS with 0.1% saponin and 2.5% normal donkey serum. Cells were then treated with primary antibodies (see list below for further information) in blocking solution overnight at 4°C. The next day, cells were washed 3x in PBS then incubated in secondary antibodies (Alexa Fluor 488, 568 and 647) at a concentration of 1:500 in blocking solution for one hour at room temperature and then HOECHST. Cells were washed in 3x in PBS and imaged on the Opera Phenix High-Content Screening System (PerkinElmer) using the Harmony analysis software.

### List of Antibodies used in this study

MBP (1:250, Abcam7349)

SOX10 (1:200, R&D Systems, AF2864)

Olig2 (1:500, EMD Millipore AB9610)

O4 (1:50, hybridoma, gift from Dr. J. Goldman)

Donkey anti-rat Alexa Fluor 555 (1:500, Invitrogen, A21434)

Donkey anti-rat Alexa Fluor 488 (1:500, Invitrogen, A21208)

Donkey anti-goat Alexa Fluor 555 (1:500, Invitrogen, A21432)

Donkey anti-rabbit Alexa Fluor 647 (1:500, Invitrogen, A31573)

Goat anti-mouse Alexa Fluor 488 IgM (1:500, Invitrogen, A21042)

Goat anti-rat Alexa Fluor 555 (1:500, Invitrogen, A21434)

Goat anti-rabbit Alexa Fluor 647 (1:500, Invitrogen, A21244)

### RNAscope and oligodendrocyte size measurements

At day 82 of the glial differentiation protocol, cells were dissociated, and replated in 96-well plates as described. For measuring oligodendrocyte size, cells were fixed in 4%PFA and stained fo MBP as described above. Images were acquired at 40X using the Opera Phenix High-Content Screening System (PerkinElmer). Harmony softwarewas used to measure the area of MBP^+^ signal around the nucleus. For RNAscope experiments, cells were fixed 2 days after plating following the protocol provided by ACDBio. In brief, cells were washed once in PBS then fixed in 10% Neutral Buffered Formalin for 30 minutes at room temperature. After 2x PBS washes, cells were dehydrated by incubation in 50% ethanol at room temperature for 5 minutes, followed by 70% ethanol at room temperature for 5 minutes, then 100% ethanol at room temperature for 5 minutes. 100% ethanol was then removed and replaced with fresh 70% ethanol at room temperature for 10 minutes. Cells were then stored at −20°C. For the RNAscope assay, cells were rehydrated by incubation in 70% ethanol (200µL/well) at room temperature for 2 minutes, followed by 50% ethanol (200μL/well) at room temperature for 2 minutes, followed by PBS at room temperature for 1 minute, then finally in PBS for 10 minutes (200μL/well). PBS was removed, plates were placed in the Humidity Control Tray provided by ACDbio, and 33uL freshly diluted Protease III (1:10 in PBS) was added in each well. Humidity Control Tray was closed and incubated for 10 minutes at room temperature. Cells were then washed in PBS for 2 minutes (200μL/well) and this was repeated twice for a total of three washes.

Staining was then performed using the ACDbio “Tech Note for using RNAscope HiPlex Alternate Display Module”, and imaging was performed using the Opera Phenix High-Content Screening System (PerkinElmer). To quantify the RNA within each cell, we used a custom-made Python script for this experiment. First, the script calculated the maximum value of the reference wells for removing the contribution from the background. Then, for each dye we computed the max projection from the stack, we subtracted the background value found above, flat field corrected each image and finally normalized them between 0 and 1. Once we had one single image for each dye, we located the cells based on the nuclei image (using an Otsu’s threshold based approach). Then we similarly located the positive portion of the images for the dyes that identified the cell type (GFAP, MBP and PLP1), and selected all the nuclei that were positive for any dye. Then looped over each of the selected cells for each of the RNA stains and inspected whether there were positive pixels within a specified region of interest. Since the size of each RNA was of about 20 pixels, we divided the number of positive pixels by 20 to get the number of total RNAs. This analysis generated a csv with one row for each of the analyzed cells, and one column for the reported number of RNAs and positivity for each of the reference dyes.

### Statistics

The Graphpad Prism software was used for all statistical analyses. The statistical test used, n, and meaning of each datapoint is described in the figure legends. The definition of center and dispersion measures is also indicated in the figure legends. Statistical significance was considered as p<0.05 (*=p<.05; **=p<.01; ***=p<.001; ****=p<.0001).

**Table 2.** iPSC lines used for each experiment.

| Line | HC 01 | HC 02 | HC 03 | HC 04 | HC 05 | RR 01 | RR 02 | RR 03 | RR 04 | RR 05 | RR 06 | SP 01 | SP 02 | SP 03 | SP 04 | SP 05 | SP 06 | PP 01 | PP 02 | PP 03 | PP 04 | PP 05 |
| --- | --- | --- | --- | --- | --- | --- | --- | --- | --- | --- | --- | --- | --- | --- | --- | --- | --- | --- | --- | --- | --- | --- |
| scRNAseq | x | x | x | x |  | x | x | x | x |  |  | x | x | x | x |  |  | x | x | x | x |  |
| RNAscope | x | x | x | x | x | x | x | x | x |  |  | x | x | x | x |  |  | x | x | x | x | x |
| Keto treatment | x |  | x | x |  | x | x | x | x | x |  | x | x | x | x | x | x |  |  | x | x | x |
| Tasin treatment | x | x | x | x | x |  |  |  |  |  |  |  | x | x | x |  |  | x |  |  | x |  |
| iPSC-cortical neurons | x | x | x | x | x | x | x | x | x | x | x | x | x | x | x | x | x | x | x | x | x | x |
| IF staining in unsorted cultures | x | x | x | x | x | x | x | x | x |  |  | x | x | x | x |  |  | x | x | x | x | x |
| OL morphology | x | x | x | x | x | x | x | x | x |  |  | x | x | x | x |  |  | x | x | x | x | x |
| IF staining for OL quantification | x | x | x | x | x | x | x | x | x |  |  | x | x | x | x |  |  | x | x | x | x | x |

---

**Figure S1.**
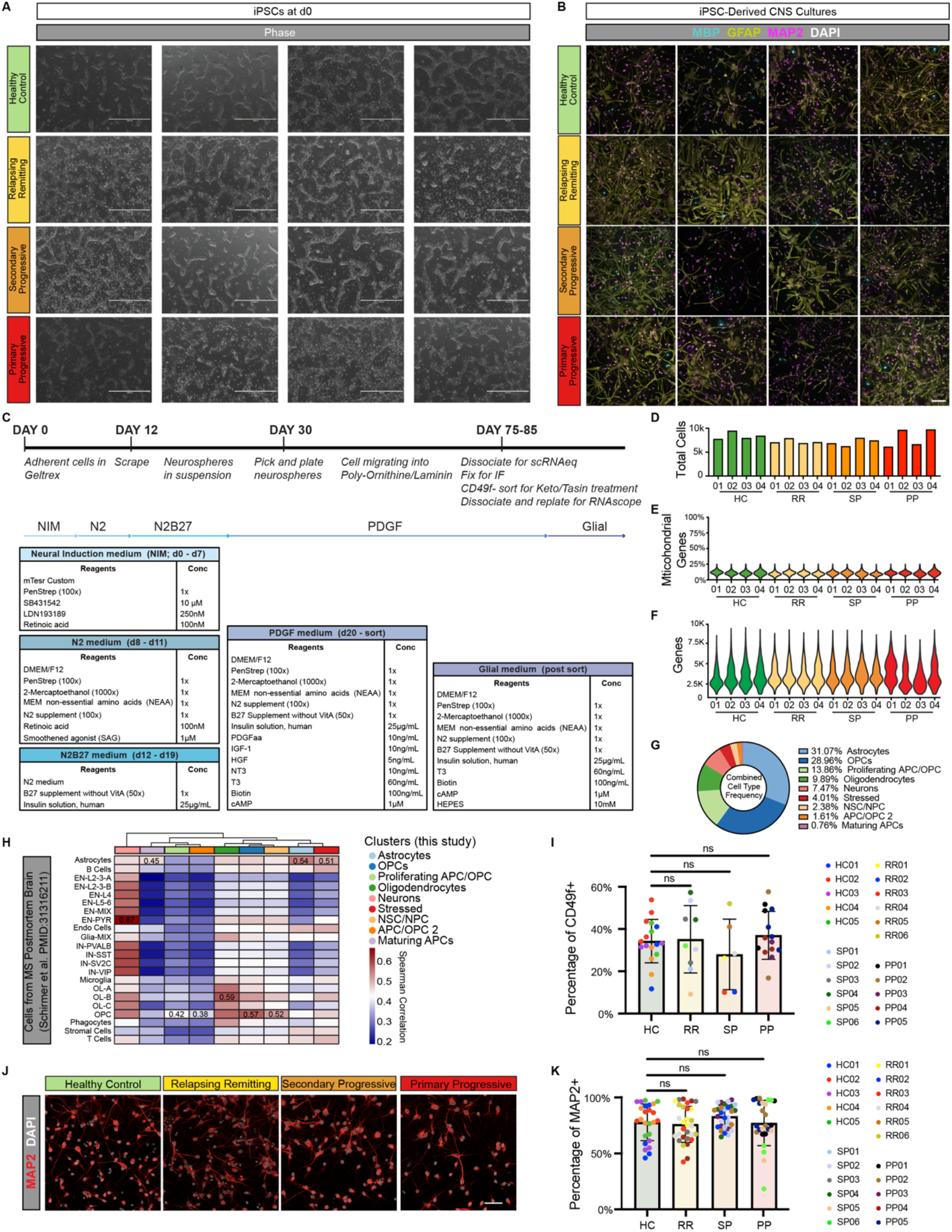
Characterization of iPSC-derived CNS cultures from people with MS. (A) Representative images of undifferentiated iPSC cultures from all 16 lines used for scRNAseq analysis. Scale bar is 1mm. (B) Representative images of iPSC-derived CNS cultures at the end of the differentiation from all 16 lines used for scRNAseq analysis. Cultures are stained for the mature oligodendrocyte marker MBP (teal), the astrocyte marker GFAP (yellow), and the neuron maker MAP2 (pink). Scale bar is 100µm. (C) Protocol and media conditions for iPSC differentiation into CNS cells. (D) Total cells per sample used for scRNAseq analysis. (E) Percentage of mitochondrial genes in each scRNAseq sample. (F) Gene counts in each scRNAseq sample. (G) Distribution of cell types from scRNAseq analysis. (H) Heatmap depicting the correlation between scRNAseq clusters identified in this study and cell types from scRNAseq analysis of MS brain tissue (Schirmer et al. PMID: 31316211). Spearman correlation values generated using the R package ClustifyR. (I) Percentage of cells in healthy control (HC), relapsing remitting (RR), secondary progressive (SP), and primary progressive (PP) iPSC-derived CNS cultures that are positive for the astrocyte marker CD49f. Data is presented as mean +/- standard deviation for *n* = 5-6 lines per group (technical replicates indicated by color-coding each line) with p-values generated by one-way Anova with Dunnett’s correction for multiple comparisons. (J) Representative images of iPSC-derived neuronal cultures stained for the neuron marker MAP2 (red). Scale bar is 50µm. (K) Percentage of cells in healthy control (HC), relapsing remitting (RR), secondary progressive (SP), and primary progressive (PP) iPSC-derived CNS cultures that are positive for the neuron marker MAP2. Data is presented as mean +/- standard deviation for *n* = 5-6 lines per group (technical replicates indicated by color-coding each line) with p-values generated by one-way ANOVA with Dunnett’s correction for multiple comparisons.

**Figure S2.**
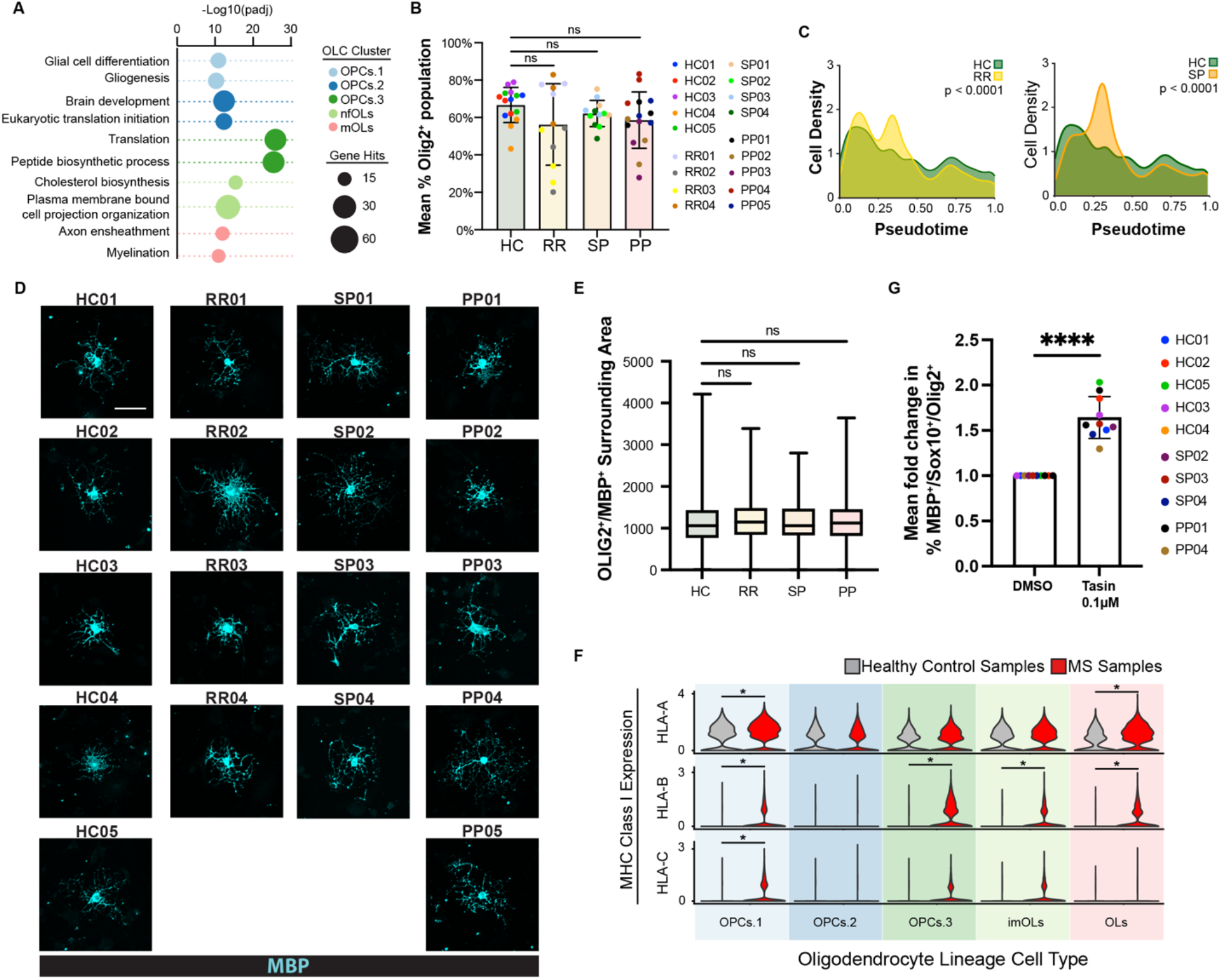
Characterization of iPSC-derived oligodendrocyte lineage cells from people with MS. (A) Gene ontology analysis of genes enriched in each oligodendrocyte lineage cell type. (B) The percentage of oligodendrocyte lineage cells in iPSC-derived cultures from healthy control (HC), relapsing remitting (RR), secondary progressive (SP), and primary progressive lines that are positive for the pan-oligodendrocyte lineage marker OLIG2. Error bars show mean ± standard deviation (n= 3 technical replicates per line for 4-5 lines per group). p-values generated by one-way ANOVA with Dunnett’s correction for multiple comparisons. (C) Cell density plot that shows the distribution of cells across the oligodendrocyte lineage trajectory from OPCs to mature oligodendrocytes for iPSC-derived cells from healthy controls (HC), relapsing remitting MS (RR), and secondary progressive MS (SP) patients. p-value generated with a two-sample Kolmogorov-Smirnov test. (D) Representative images of mature oligodendrocytes from healthy control (HC), relapsing remitting (RR), secondary progressive (SP), and primary progressive cultures stained with the mature oligodendrocyte marker MBP. Scale bar is 50µm. (E) Quantification of MBP^+^ oligodendrocyte area for oligodendrocytes from healthy control (HC), relapsing remitting (RR), secondary progressive (SP), and primary progressive cultures. Data is presented as mean +/- standard deviation for *n* = 299; 150; 254; 234 cells, per respective group. p-values generated by one-way ANOVA with Dunnett’s correction for multiple comparisons. (F) Expression of MHC Class I genes in oligodendrocyte lineage cell types from healthy control and MS cultures. p-value generated by Wilcoxon ranked sum test within the Seurat R package. * p < 0.05. (G) Fold-change in the percentage of MBP^+^ cells in all cultures treated with vehicle (DMSO) or the oligodendrocyte enhancing compound TASIN-1 at 100nM. Data is presented as mean +/- standard deviation for *n* = 10 lines. p-values generated by two-way unpaired t-test.

**Figure S3.**
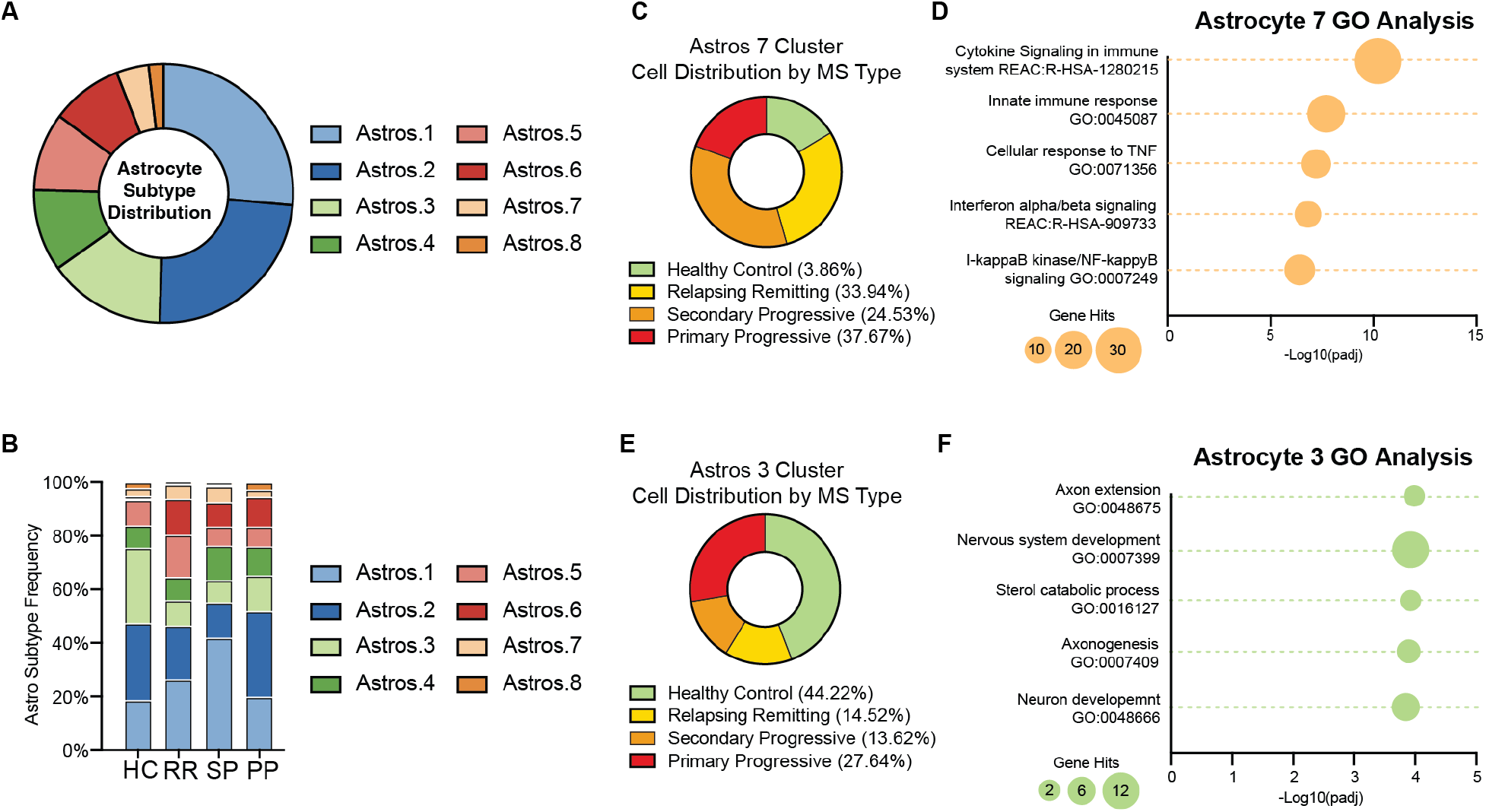
Reactive astrocyte subtypes in iPSC-derived astrocytes from people with MS. (A) The proportion of each astrocyte subtype in iPSC-derived cultures from all samples. (B) The proportion of each astrocyte subtype in iPSC-derived cultures from healthy control (HC), relapsing remitting (RR), secondary progressive (SP), and primary progressive lines. Astrocyte subclusters 6 and 7 are enriched in cultures from people with MS while astrocyte subcluster 3 is depleted in cultures from people with MS. (C) Distribution of healthy control, relapsing remitting, secondary progressive, and primary progressive iPSC-derived astrocytes in the astrocyte subcluster 7. (D) Gene ontology analysis of genes significantly increased in astrocytes subcluster 7 compared to all other astrocyte subclusters. (E) Distribution of healthy control, relapsing remitting, secondary progressive, and primary progressive iPSC-derived astrocytes in the astrocyte subcluster 3. (F) Gene ontology analysis of genes significantly increased in astrocytes subcluster 3 compared to all other astrocyte subclusters.

**Figure S4.**
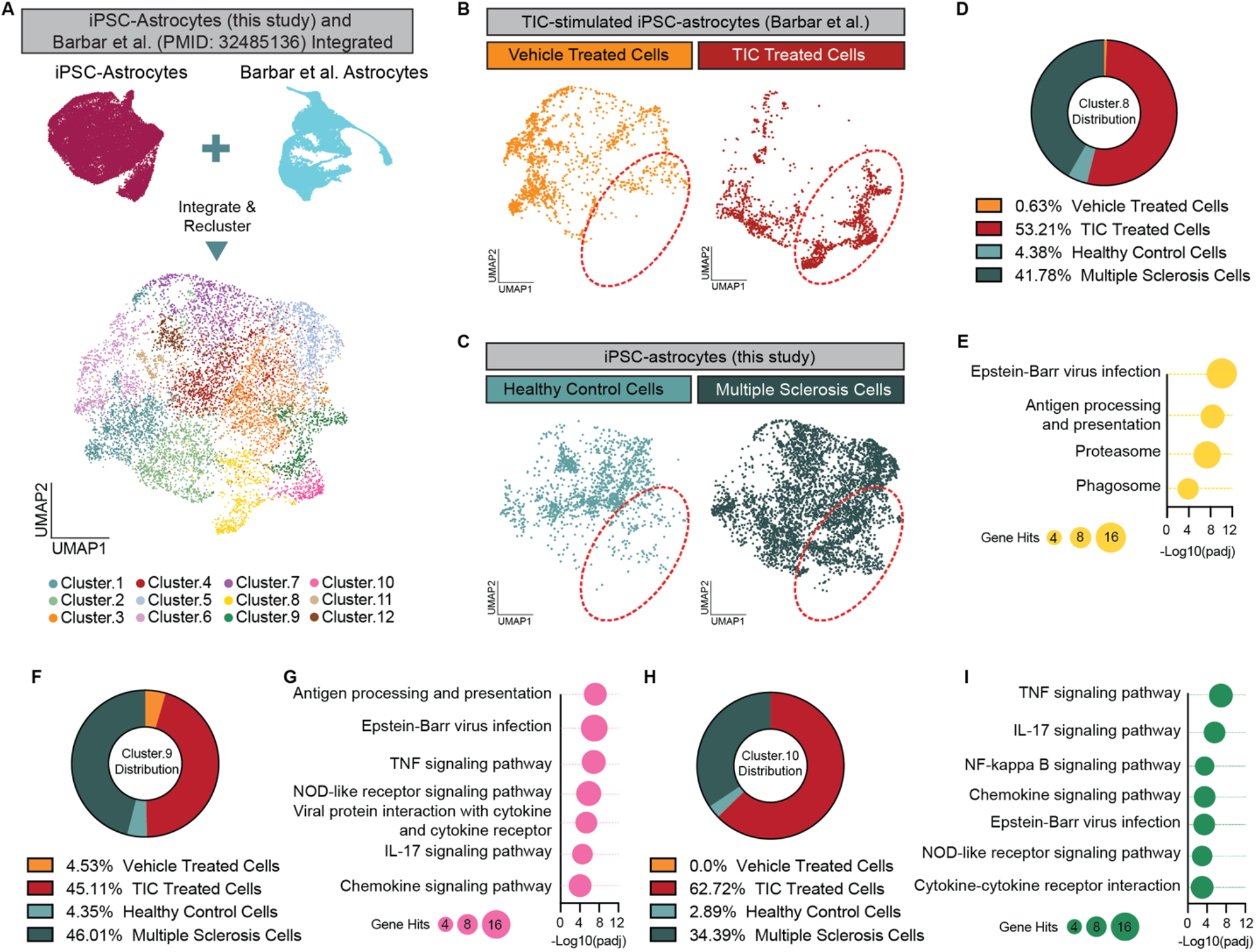
iPSC-derived astrocytes from people with MS share a transcriptional profile with inflammatory-driven human iPSC-derived reactive astrocytes. (A) UMAP plot of astrocytes from this study integrated with iPSC-derived astrocytes from healthy control individuals exposed to either vehicle, or the inflammatory cytokines TNF, IL1α, and C1q (TIC) (Barbar et al. PMID: 32485136). (B) UMAP plots showing the distribution of cells from vehicle-treated and TIC-treated iPSC-derived astrocytes. Red circle highlights Clusters 8, 9, and 10 which are enriched for iPSC-derived astrocytes exposed to TNF, IL1α, and C1q (TIC). (C) UMAP plots showing the distribution of cells from iPSC-derived astrocytes from healthy control individuals and individuals with MS. Red circle highlights Clusters 8, 9, and 10 which are enriched for iPSC-derived astrocytes from people with MS. (D) Distribution of vehicle-treated iPSC-derived astrocytes (yellow), TIC-treated iPSC-derived astrocytes (red), healthy control iPSC-derived astrocytes (light blue), and MS iPSC-derived astrocytes (dark blue) in Cluster 8 of the integrated data sets. (E) Gene ontology analysis of the top 100 genes enriched in Cluster 8 of the integrated data set compared to all other clusters in the integrated data set. (F) Distribution of vehicle-treated iPSC-derived astrocytes (yellow), TIC-treated iPSC-derived astrocytes (red), healthy control iPSC-derived astrocytes (light blue), and MS iPSC-derived astrocytes (dark blue) in Cluster 9 of the integrated data sets. (G) Gene ontology analysis of the top 100 genes enriched in Cluster 9 of the integrated data set compared to all other clusters in the integrated data set. (H) Distribution of vehicle-treated iPSC-derived astrocytes (yellow), TIC-treated iPSC-derived astrocytes (red), healthy control iPSC-derived astrocytes (light blue), and MS iPSC-derived astrocytes (dark blue) in Cluster 10 of the integrated data sets. (I) Gene ontology analysis of the top 100 genes enriched in Cluster 10 of the integrated data set compared to all other clusters in the integrated data set.

